# Hidden biotic stress alters climate sensitivity in woody plants

**DOI:** 10.64898/2026.07.18.739329

**Authors:** Eduardo Moralejo, Marina Montesinos, Blanca Landa, Diego Olmo

## Abstract

Chronic infection by vascular pathogens is conventionally expected to severely constrain host biomass accumulation, yet empirical evidence from long-lived woody plants remains inconsistent. We investigated the long-term impacts of *Xylella fastidiosa* colonization on the radial growth and climate sensitivity of adult Mediterranean almond trees (*Prunus dulcis*), aiming to resolve how persistent vascular infections modulate tree performance and resilience under a changing climate. We coupled high-resolution dendrochronological analysis with a novel, ring-resolved molecular reconstruction of individual infection histories across 706 annual rings. This hindcasting approach allowed for the retrospective identification of precise colonization dates, bacterial loads (Ct values), and pathogen subspecies (subsp. *fastidiosa* vs. subsp. *multiplex*). Growth–climate relationships were modelled using standardized ring-width indices (RWI) against a crop-weighted water deficit index (CWDi). Intra-host colonization followed a steep radial gradient, with active bacterial abundance concentrated in newly formed, functional outer xylem. Surprisingly, chronic infection did not trigger a sustained reduction in baseline annual ring width. Instead, pathogens fundamentally reshaped climate–growth sensitivity. Hosts infected by subsp. *fastidiosa* maintained high plastic growth tracking during wet years, whereas this capacity was significantly attenuated in those harbouring subsp. *multiplex*. Despite the absence of a chronic signal in trunk radial growth, vascular impairment was tightly associated with severe retrograde canopy dieback. Our findings indicate that chronic infection by *X. fastidiosa* can act as a latent biotic stressor, altering host physiological sensitivity to environmental fluctuations without directly suppressing baseline stem growth. This pattern is consistent with the marked temporal decoupling between spring cambial activity and late-summer bacterial proliferation, together with progressive vascular dysfunction leading to severe canopy dieback. These results suggest that current abiotic-centred frameworks of drought-induced decline may underestimate the contribution of cryptic vascular pathogens to vegetation mortality under intensifying climate change.

## Introduction

Global change is accelerating the frequency, intensity, and duration of droughts worldwide, with profound consequences for woody vegetation, forest productivity, and ecosystem stability (Carnicer *et al*., 2011; Choat *et al*., 2012; Torres-Ruiz *et al*., 2024). Across biomes, recent episodes of tree decline and mortality are commonly interpreted through the framework of drought-induced hydraulic failure. Under this paradigm, declining soil moisture elevates xylem tension and promotes cavitation, driving a progressive loss of hydraulic conductivity until critical thresholds are breached (Anderegg *et al*., 2015a, 2016; Venturas *et al*., 2017). This framework has become central to predicting vegetation vulnerability under climate change and to explaining large-scale mortality events in both forests and orchards (Sanchez-Martinez *et al*., 2023).

Tree performance under drought is, however, rarely shaped by climate alone. Pathogens and pests are conventionally treated as secondary consequences of water stress, linked to drought-induced susceptibility, opportunistic invasion, or post-stress decline (Desprez-Loustau *et al*., 2006; Oliva *et al*., 2014; Anderegg *et al*., 2015b; Teshome *et al*., 2020). Concurrently, many vascular microorganisms persist within host xylem for years, potentially altering plant function long before severe drought manifests (Bettenfeld *et al*., 2020; Anguita-Maeso *et al*., 2022).

Despite this, xylem-inhabiting microorganisms have been studied primarily through the lens of agricultural disease (Yadeta & J. Thomma, 2013; De La Fuente *et al*., 2022), leaving their broader ecological roles in natural ecosystems poorly understood (Morris *et al*., 2009; García-Guzmán & Heil, 2014). This knowledge gap is increasingly addressed by viewing plant– microbe interactions as a continuum of symbiosis, where microbial taxa shift between mutualistic, commensal, or pathogenic states depending on environmental conditions, host associations, and life-history stages (Casadevall & Pirofski, 2003; Morris *et al*., 2022; Stengel *et al*., 2022).

Among vascular microorganisms displaying such plastic lifestyles, *Xylella fastidiosa* provides an ideal model system. This xylem-limited, vector-borne bacterium colonizes vessels, forms biofilms, and induces host defensive responses—such as tyloses and gels—that restrict water transport across a broad range of woody hosts (De La Fuente *et al*., 2022). Disease symptoms, including leaf scorch, branch dieback, and mortality, strongly overlap with phenotypes attributed to severe water stress (Almeida & Nunney, 2015). Crucially, many infected hosts remain asymptomatic for extended periods, suggesting that vascular impairment long precedes visible decline, which likely leads to an underestimation of pathogen prevalence in woody communities (Chatterjee *et al*., 2008; Morris & Moury, 2019). Although *X. fastidiosa* is native to the Neotropics and North America (Sicard *et al*., 2018) —where it is believe to persist as a largely asymptomatic endophyte (Newman *et al*., 2004; Almeida & Nunney, 2015) —its ecological role in natural forest ecosystems remains elusive, as most knowledge derives from agricultural invasions (Morris & Moury, 2019). Yet, its likely ancient evolutionary history (Morris & Moury, 2019), coupled with the high diversity of xylem-feeding vectors in tropical forests (Feng *et al*., 2024), points to long-term ecological associations within these complex ecosystems. Furthermore, the recent identification of sister *Xylella* species in Asia suggests that the global diversity and prevalence of this genus remain substantially underestimated (Briand *et al*., 2025).

Recent work has begun to explicitly merge plant hydraulics with the vascular pathology of *X. fastidiosa* (Sabella *et al*., 2019; Walker *et al*., 2024). Experimental and conceptual studies indicate that conduit anatomy and vessel dimensions influence host vulnerability to vascular colonization, while the pathogen itself exacerbates hydraulic dysfunction (Montilon *et al*., 2023). In grapevine and olive, for instance, xylem anatomical traits linked to embolism resistance may also confer tolerance against *X. fastidiosa* (Fry & Milholland, 1990; Carluccio *et al*., 2023). Together, these findings demonstrate that drought stress and vascular infection are not independent pressures, but interacting forces operating on the same transport system.

Nevertheless, whether chronic vascular infection consistently reduces tree growth remains unresolved. Direct suppression may be weak, transient, or offset by host tolerance; instead, pathogen impacts may manifest primarily as altered sensitivity to climatic variability.

Distinguishing between these alternatives requires long-term retrospective approaches capable of resolving host performance before and after infection. Tree-ring analysis offers a powerful but underutilized framework for this purpose, because annual radial growth integrates carbon balance, hydraulic function, and cumulative stress exposure over time (Brzostek *et al*., 2014). Growth chronologies can thus reveal whether vascular pathogens reduce baseline vigour, amplify drought sensitivity, or trigger delayed declines preceding visible collapse (Cailleret *et al*., 2017).

Here, we use almond trees (*Prunus dulcis*), a drought-exposed Mediterranean crop highly susceptible to *X. fastidiosa*, as a natural model system (see Supplementary note). In Mallorca, almond leaf scorch disease (ALSD) has affected more than one million trees and is associated with the subspecies *fastidiosa* and *multiplex*, mirroring the situation in California (Chen *et al*., 2005; Moralejo *et al*., 2020). To capture this landscape-scale phenomenon, we developed tree-ring chronologies from a representative subset of 30 trees, selected from a broader population of hundreds of thousands of individuals sharing the same symptom development pattern. For these trees, infection status, bacterial load, subspecies identity, and climatic covariates were fully characterized. We investigated whether chronic vascular infection leaves a detectable imprint on radial growth and whether it alters the relationship between tree growth and climatic variability. By integrating tree-ring ecology with plant pathology, this study provides evidence that vascular pathogens may shape woody plant performance under climate stress.

## Materials and Methods

### Study system and sampling

The study was conducted in rainfed almond orchards across Mallorca (Balearic Islands, Spain), a Mediterranean region characterized by high interannual precipitation variability. Almond orchards in this area are cultivated under semi-arid conditions, with mean annual rainfall ranging from 300 to 600 mm. More than 89 local almond cultivars have been characterized on the island; yet, despite this high genetic and varietal diversity, the ALSD epidemic currently affects over 85% of all almond trees, exhibiting a consistent symptom progression across the landscape.

During the *X. fastidiosa* outbreak, European Union emergency phytosanitary regulations mandated the eradication of all host trees testing positive for the pathogen (Quetglas *et al*., 2022). Seizing this regulatory context, trunk cross-sections were collected from infected almond trees felled between 2017 and 2018 during official eradication campaigns. All sampled individuals had previously tested positive for *X. fastidiosa* via official diagnostic protocols. For each tree, geographic coordinates were recorded. Three stem cross-sections ∼10 cm thick, including one at breast height, 1.3 m, and two main branches were harvested using a chainsaw and stored in labelled bags for laboratory analysis. Although 34 infected trees were initially sampled, 30 individuals yielded cross-sections with sufficient anatomical quality to build reliable tree-ring chronologies, forming the core dataset characterized for infection status, bacterial load, and climatic covariates.

### Wood preparation and tree-ring analyses

In the laboratory, wood sections were surfaced using an electric planer and progressively sanded with increasingly fine grit sandpaper (up to 600-grit) until cell walls were clearly visible under magnification. Annual growth rings were visually identified under a stereomicroscope and cross-dated following standard dendrochronological procedures. Ring boundaries along one to three radial transects per cross-section were carefully marked with a scalpel before measurement. Ring widths were then measured to the nearest 0.01 mm along these transects using high-resolution scanned images (minimum 1200 dpi) analysed with the calibrated image-analysis software ImageJ (Fig. 1).

**Figure 1.**
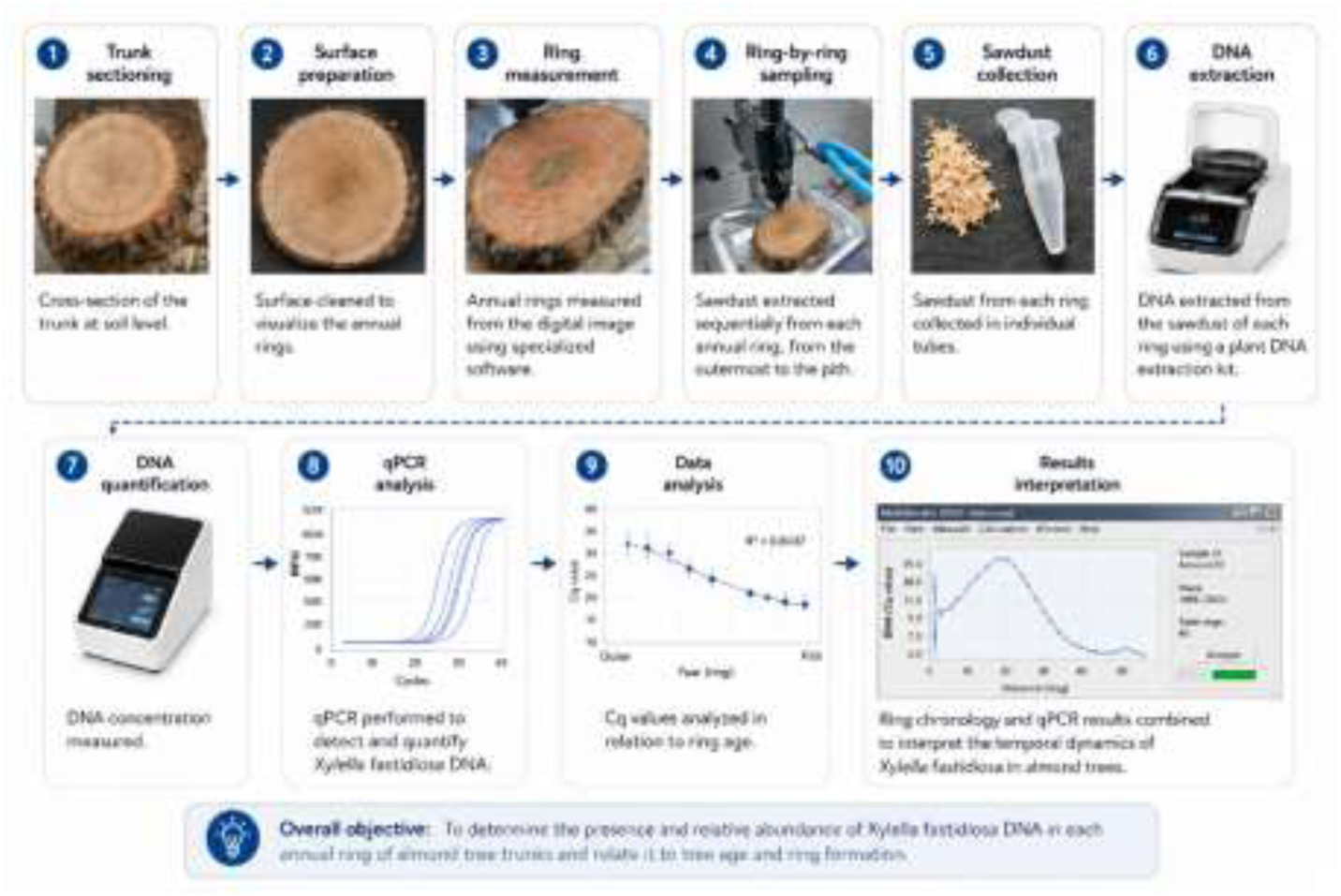
Methodological workflow for ring-resolved dendrochronological and molecular analysis. The ten-step pipeline illustrates the retrospective tracking of pathogen dynamics: (1) trunk cross-sectioning at breast height; (2) surface planing and sanding for ring visualization; (3) digital ring measurement and cross-dating via specialized software; (4) sequential ring-by- ring micro-sampling from the outermost ring to the pith; (5) wood dust collection into individual sterile tubes; (6) genomic DNA extraction; (7) DNA quantification; (8) target qPCR amplification for *X. fastidiosa*; (9) analysis of cycle quantification (Ct) values across the temporal pith-to-bark gradient; and (10) integration of tree-ring chronologies and molecular data to reconstruct long-term epidemic dynamics.

Cross-dating quality was evaluated both visually and statistically using the dplR package in R (Bunn, 2008). Series displaying ambiguous ring boundaries or poor statistical alignment were excluded from subsequent analyses, resulting in the final selection of 30 trees. To remove age-related biological growth trends and minimize size-related variances among trees, raw ring-width series were detrended and standardized into dimensionless ring-width indices (RWI).

### Conceptual model of bacterial distribution across growth rings

Our interpretation of bacterial DNA abundance across successive growth rings was based on the known anatomy and development of almond xylem. Each year, the vascular cambium produces a new ring of functional xylem that becomes hydraulically connected to newly formed shoots and leaves. Because *X. fastidiosa* is transmitted to and proliferates within active xylem vessels, we assumed that newly formed outer rings have a greater probability of receiving bacterial inoculum and sustaining bacterial multiplication than older inner rings. Consequently, bacterial DNA was expected to be concentrated in the most recently formed xylem and to decrease progressively toward older rings as hydraulic activity declines and connections with contemporary foliage are lost. Under this conceptual framework, Ct values were predicted to increase from outer to inner rings, reflecting a decline in bacterial abundance with increasing ring age (Fig. 2). To test this prediction, Ct values obtained from annual growth rings were analysed as a function of ring age (distance from the outermost ring), pathogen subspecies, and their interaction.

**Figure 2.**
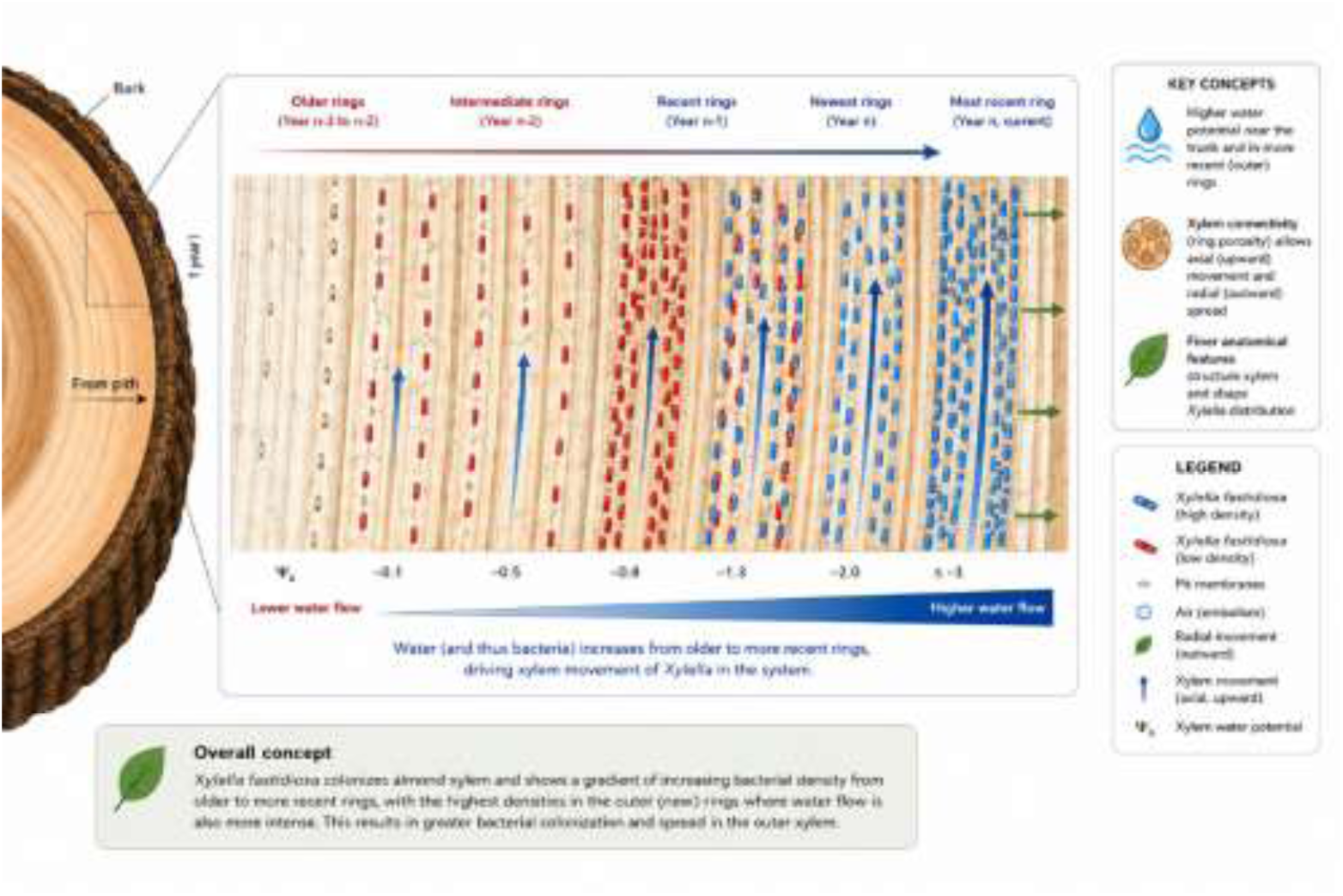
Conceptual model of *Xylella fastidiosa* persistence and movement across annual rings in almond trunks. The figure illustrates the proposed spatial dynamics of *X. fastidiosa* within the xylem of almond tree trunks. Water transport and hydraulic conductivity are highest in the most recent outer rings, promoting bacterial survival, multiplication and passive movement through xylem vessels and pit membrane connections. Live bacterial cells (blue) are concentrated mainly in the current and recently formed rings, whereas dead bacterial cells (red) accumulate progressively in older rings toward the pith. Radial movement within rings and limited axial movement between adjacent rings are facilitated by xylem connectivity and sap flow. Older inner rings progressively lose hydraulic activity and become largely unsuitable for bacterial persistence, resulting in the absence of detectable cells in the oldest heartwood regions. The model highlights how xylem age, hydraulic function and ring connectivity may structure the long-term persistence of *X. fastidiosa* in perennial woody hosts.

### Ring-resolved molecular reconstruction of infection history

To reconstruct the temporal progression of *X. fastidiosa* colonization within individual hosts, the outermost 20 annual growth rings of each stem section were micro-sampled for molecular analysis. Annual rings were visually identified under a stereomicroscope, and approximately 0.5 mg of wood dust was extracted from each individual ring using a vertical drill stand fitted with a 0.4-mm drill bit. Because annual ring widths occasionally approached the diameter of the drill bit, micro-sampling could inadvertently include a small amount of tissue from an adjacent ring. Accordingly, infection dates inferred from ring-specific qPCR analyses were interpreted with an estimated temporal uncertainty of ±1 year. To prevent cross-contamination, the initial wood particles generated at the surface of each drilling site were discarded. Rings were sampled sequentially along the pith-to-bark gradient (from older to younger wood). Between consecutive ring extractions, all drilling equipment and sample surfaces were thoroughly disinfected with 70% ethanol. Wood dust from each ring was collected into sterile bags and homogenized using a Homex 6 grinder (Bioreba Instruments, Switzerland).

DNA extracted from the ring-derived wood dust was analysed via quantitative PCR (qPCR) in duplicate at both the Official Plant Health Laboratory of the Balearic Islands (LOSVIB) and the IAS-CSIC facilities. Cycle quantification (Ct) values served as a proxy for bacterial DNA abundance across the temporal profile of the stem. Positive detections (Ct < 37) were utilized to infer the approximate onset year of vascular infection for each tree and to reconstruct the chronology of disease establishment at the population level. This hindcasting approach assumes that while active xylem transport and viable bacterial populations are concentrated in the most recently formed, functional outermost rings, residual pathogen DNA remains entrapped and persists chronologically within the older, non-functional wood matrix (Fig. 1).

### Molecular detection and subspecies identification of *Xylella fastidiosa*

Total genomic DNA was extracted from homogenized wood tissues using the E.Z.N.A. HP Plant Mini Kit (Omega Bio-Tek, Norcross, GA, USA), strictly adhering to the European and Mediterranean Plant Protection Organization (EPPO) diagnostic standards. Diagnostic detection of *X. fastidiosa* was performed using two complementary species-specific qPCR assays: (i) the Harper *et al*., (2010) using primers XF-F/XF-R and probe XF-P, and (ii) the protocol of Francis *et al*., (2006) based on primers HL5/HL6 and their corresponding TaqMan probe. All real-time amplification reactions were executed in an Eco Real-Time PCR System (PCRmax, Staffordshire, UK).

Pathogen subspecies assignment was conducted on a representative subset of positive samples using a multiplex duplex TaqMan qPCR assay designed to discriminate between the subspecies *fastidiosa* and *multiplex* (Dupas *et al*., 2019). To further resolve the genetic lineage at the strain level, multilocus sequence typing (MLST) was performed by amplifying and sequencing seven standard housekeeping genes (*leuA, petC, malF, cysG, gltT, holC*, and *nuoL*). Sequence Types (STs) were determined by aligning consensus sequences against the public *X. fastidiosa* MLST database (PubMLST).

### Climatic data and climatic water deficit

To evaluate whether water stress in almond trees was driven by climatic conditions, we analysed long-term climatic data from Mallorca. Monthly temperature and precipitation records from nine meteorological stations distributed across the island were obtained from the Spanish Meteorological Agency (AEMET) for the period 1988–2017. Monthly potential evapotranspiration (ET₀) was estimated using the Hargreaves equation:

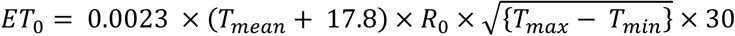

where *T_mean_*, *T_max_*, and *T_min_* represent the monthly mean, maximum, and minimum temperatures (°C), respectively, and *R_0_* is the extra-terrestrial radiation expressed as equivalent evaporation (mm d^-1^). To quantify cumulative water stress, we calculated a crop-weighted water deficit index (CWDi, mm) annually over the almond growing season (February–August) as follows:

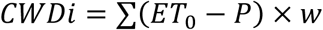

where *P* is the monthly precipitation and w is a monthly weighting factor accounting for seasonal canopy development. Based on regional phenology, the monthly weights (*w*) were set to 0.05, 0.10, 0.20, 0.30, 0.20, 0.10, and 0.05 from February to August, respectively.

### Infection history and epidemiological variables

Infection dynamics were operationalized on a tree-year basis using the chronologically reconstructed onset dates of infection. Each annual growth ring observation was treated as a binary state, classified as either non-infected (0) or infected (1). For trees with confirmed positive detections, the initial year of inferred colonization established the baseline to calculate a time-series variable representing years since infection for all subsequent annual rings. To characterize terminal disease expression, visual canopy symptom severity was recorded during the final 2017 field assessment using a semi-quantitative 6-class ordinal scale (ranging from 0 = asymptomatic canopy to 5 = dead tree; see Fig. S1 for detailed rating metrics). Individuals with unresolved or ambiguous infection timing were retained exclusively for baseline population-level growth characterizations but were systematically excluded from downstream statistical modelling that required explicit temporal infection covariates.

## Statistical analyses

Radial growth responses were analysed using linear mixed-effects models (LMMs) to evaluate the impacts of environmental regimes and disease progression on the ring width index (RWI). First, a baseline global LMM was fitted to assess macroclimatic and broad pathological constraints, specifying binary infection status (healthy vs. infected) and mean climatic water deficit (CWDi) as fixed effects. To account for the hierarchical structure of the dendrochronological data and repeated measurements within individual hosts, tree identity was incorporated as a random intercept across all mixed models. To resolve specific pathological mechanisms within the infected population, subsequent LMMs evaluated the internal pathogen dynamics. The temporal progression of bacterial colonization was modelled using chronological years as a continuous fixed predictor against Cycle threshold (Ct) values derived from qPCR.

To evaluate whether specific genetic lineages dictate differential drought sensitivity, an explicit interaction model was built specifying pathogen subspecies (*X. fastidiosa* subsp. *fastidiosa* vs. subsp. *multiplex*) and CWDi as interacting fixed factors, while adjusting for continuous bacterial load (Ct values). Post-hoc conditional distance slopes and trends for each lineage along the drought intensity gradient were estimated using estimated marginal trends. Final parameters for all LMMs were estimated via Restricted Maximum Likelihood (REML), and degrees of freedom were approximated using Satterthwaite’s method. Model assumptions, including homoscedasticity and normality of residuals, were rigorously verified through visual inspection of diagnostic plots, with statistical significance established at α = 0.05. Epidemiological dynamics and the temporal trajectory of host mortality risk were evaluated using non-parametric Kaplan–Meier survival estimators based on the chronologically reconstructed onset years of infection. Because the sampled individuals were standing and functional prior to being felled during the emergency eradication campaigns, tree life-histories were treated as right-censored observations. All statistical computations and modelling were performed using R software (v4.4.1), utilizing the *lme4* package for mixed-effects structures (Bates *et al*., 2015), *lmerTest* for hypothesis testing (Kuznetsova *et al*., 2017), *emmeans* (Lenth *et al*., 2019) and *ggeffe* (Lüdecke, 2018) Ct*s* for post-hoc marginal effects, and the *survival* and *ggsurvfit* packages for time-to-event analyses (Therneau, 2015).

## Results

### Within-host pathogen dynamics

A total of 706 annual ring observations from 30 almond trees were analysed to evaluate the effects of infection status, climatic water deficit, pathogen load, canopy symptom severity, and pathogen subspecies on detrended radial growth (RWI). Mean climatic water deficit (CWDi), representing the baseline macroclimatic forcing on almond growth, remained stable over the 30- year study period despite substantial interannual variability (Fig. 3). Against this stable climatic background, the probability of infection scaled up over time, closely mirroring the historical progression of the ALSD epidemic in Mallorca (Fig. 3 & Supplementary Material) (Moralejo *et al*., 2020).

**Figure 3.**
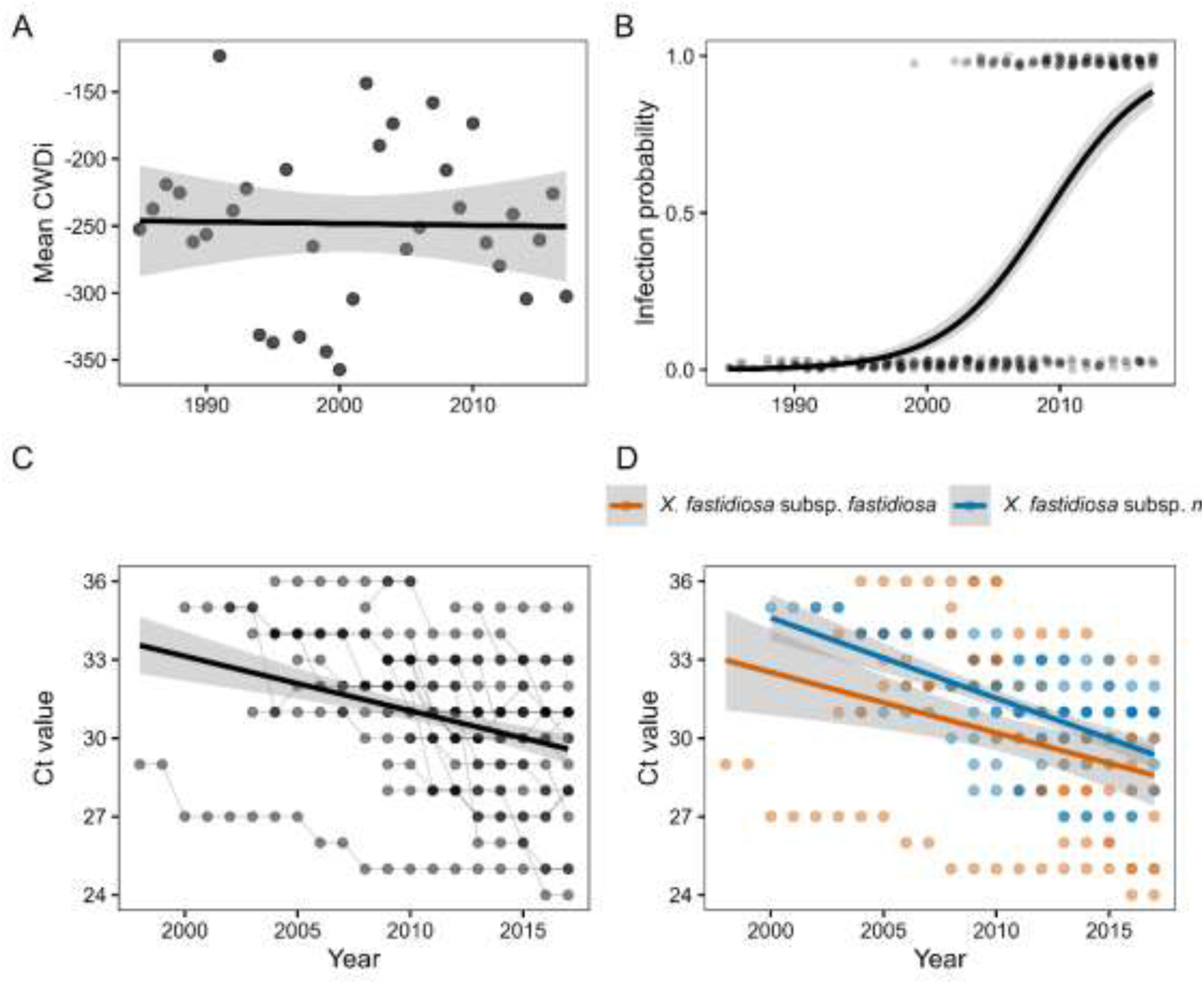
Temporal dynamics of climatic conditions, infection probability, and bacterial load in almond trees. (A) Mean annual climatic water deficit (CWDi), showing no directional temporal trend. (B) Probability of infection by *X. fastidiosa*, which increased markedly over time. (C) Ct values across infected tree rings, indicating a progressive increase in bacterial load. Thin lines represent individual trees and the solid line shows the overall trend. (D) Ct dynamics by *X. fastidiosa* subspecies. Points represent individual observations and shaded areas denote 95% confidence intervals.

Quantitative PCR analyses across successive growth rings revealed a steep radial gradient in bacterial DNA abundance. Ct values increased significantly from the outer to the inner rings (*F* = 292.9, *P* < 0.001), demonstrating substantially higher bacterial loads in recently formed, functional xylem tissues (Fig. 3). This spatial pattern, however, was highly subspecies-specific (Ring × Subspecies interaction: *F* = 6.0, *P* = 0.0152); subsp. *fastidiosa* exhibited a significantly steeper radial decline in bacterial DNA toward the pith than subsp. *multiplex*, consistent with a stricter confinement to newly active xylem vessels (Fig. 3A–D).

### Pathogen impacts are mediated through climate–growth sensitivity

Across all monitored hosts, radial growth was strongly governed by climate. In the global linear mixed-effects model assessing the combined effects of infection status and climate, CWDi exerted a dominant, significant negative effect on RWI (*F* = 59.30, *P* < 0.001; indicating that growth was strongly limited by water deficit), whereas binary infection status per se exerted no detectable main effect (*F* = 0.52, *P* = 0.47). Thus, interannual climatic variability explained the vast majority of annual growth fluctuations, while infected and non-infected individuals maintained identical baseline growth rates.

Models incorporating the high-resolution molecular estimates of bacterial load yielded consistent results, showing no evidence of direct growth suppression driven by infection intensity. In the model partitioning Ct, climate, and pathogen subspecies, bacterial load was not significantly associated with RWI (*P* = 0.67), whereas CWDi remained a powerful predictor of radial investment (*t* = 2.93, *P* < 0.004). Among infected trees with resolved pathogen identity, mean radial growth did not differ between hosts harbouring subsp. *fastidiosa* and those with subsp. *multiplex* (*t* = 0.71, *P* = 0.480), a pattern that remained unchanged after statistically controlling for internal bacterial load (*t* = 0.76, *P* = 0.448).

Concurrently, while mean growth rates were uniform across subspecies, their strategic responses to climatic fluctuations diverged sharply. The model testing the interaction between pathogen identity and climate revealed a highly significant Subspecies × CWDi interaction (*t* = - 2.72, *P* = 0.007). Trees infected by subsp. *fastidiosa* retained a plastic and strong growth response to favourable climatic conditions (moist years) (slope = 0.156 ± 0.039, *P* < 0.001), whereas this climate-tracking capacity was substantially attenuated in trees infected by subsp. *multiplex* (Fig. 4). These findings indicate that pathogen identity modulates how host growth tracks environmental variability, even in the absence of macroscopic differences in cumulative radial biomass.

**Figure 4.**
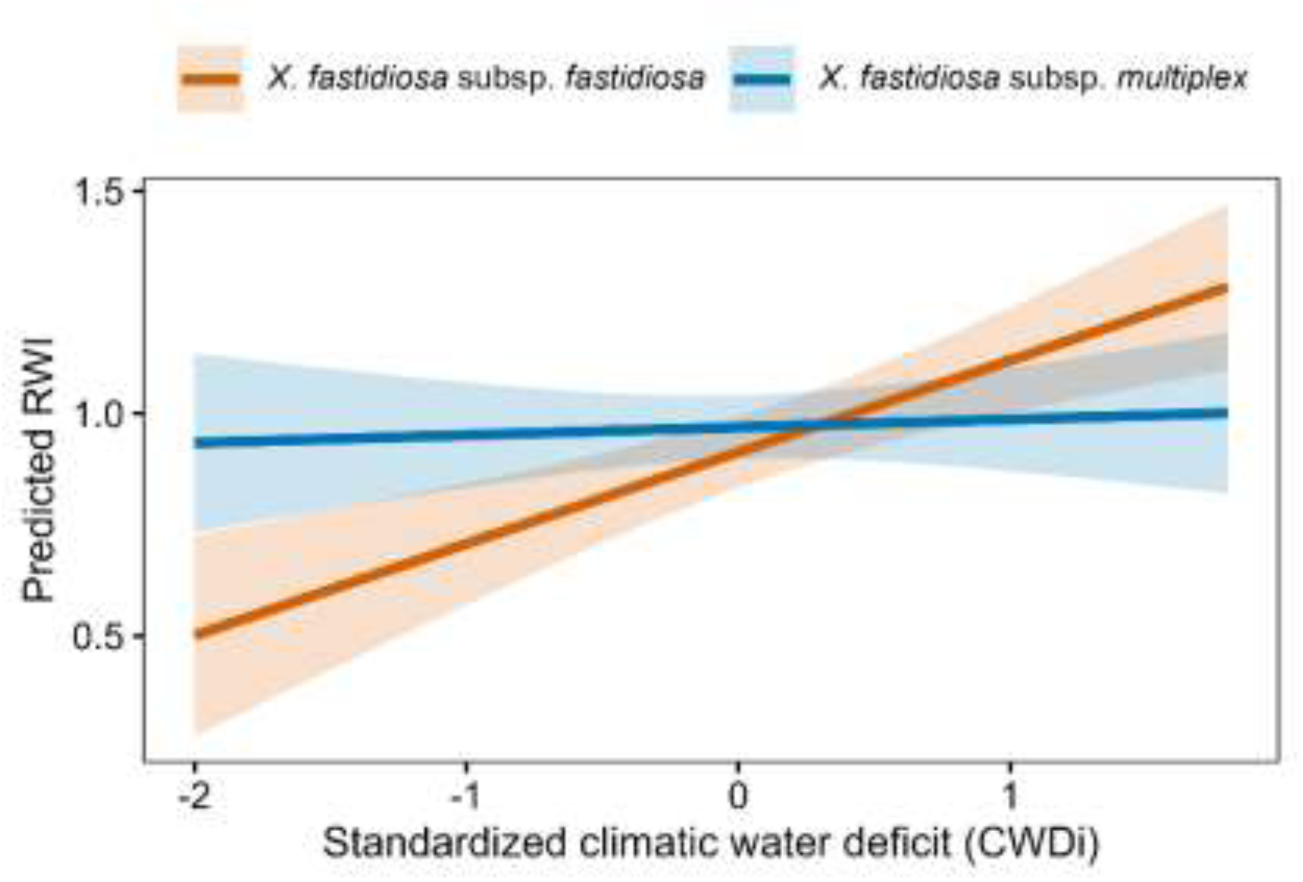
Pathogen subspecies differentially modify growth responses to climatic water deficit. Predicted radial growth (ring-width index, RWI) as a function of climatic water deficit (scaled CWDi) for almond trees infected by *Xylella fastidiosa* subsp. *fastidiosa* and subsp. *multiplex*. Shaded areas represent 95% confidence intervals. While mean growth did not differ between subspecies, their responses to climatic variability diverged markedly. Trees infected by subsp. *fastidiosa* showed a stronger positive growth response to favourable climatic conditions, whereas those infected by subsp. *multiplex* exhibited a weaker and nearly flat response. These results indicate that pathogen identity alters growth–climate sensitivity without affecting baseline growth, supporting the view that chronic infection influences host performance primarily through host–environment interactions.

### Independence of symptom severity, pathogen load, and mortality risk

None of the sampled trees exhibited signs of bark borer insect activity prior to felling, excluding secondary pest interference. Visual canopy severity scores (1–5 scale) recorded in 2017 were uncoupled from radial growth in cross-sectional analyses. In the 2017 model, variance in RWI was not explained by symptom severity (*F* = 0.39, *P* = 0.542), bacterial load (*F* = 0.16, *P* = 0.695), or subspecies identity (*F* = 0.04, *P* = 0.845). Likewise, long-term retrospective models detected no significant chronic effect of visual severity on historical growth (*P* = 0.519).

Ordinal regression models confirmed the absence of a relationship between final canopy defoliation/scorch scores and the Ct values measured in the outermost, functional ring. Furthermore, symptom severity did not differ systematically between the two pathogen subspecies.

This disease progression unfolded within a broader epidemiological context characterized by a seasonal pattern of symptom expression (Fig S2) and a significant increase in almond tree mortality recorded from 2012 to 2017 (Fig S3). Against this backdrop, the non-parametric Kaplan–Meier survival analysis revealed a progressive decline in host survival probability following the estimated onset of infection (Figure 5). The overall median survival time was estimated at 14 years post-infection (95% CI = 11–16 years; Fig S5). This timeline represents a highly conservative estimate of host longevity under pathogen pressure, as the standing, functional individuals were management-censored (felled) during mandatory emergency eradication campaigns, meaning their biological time-to-mortality could potentially extend further under unmanaged conditions.

**Figure 5.**
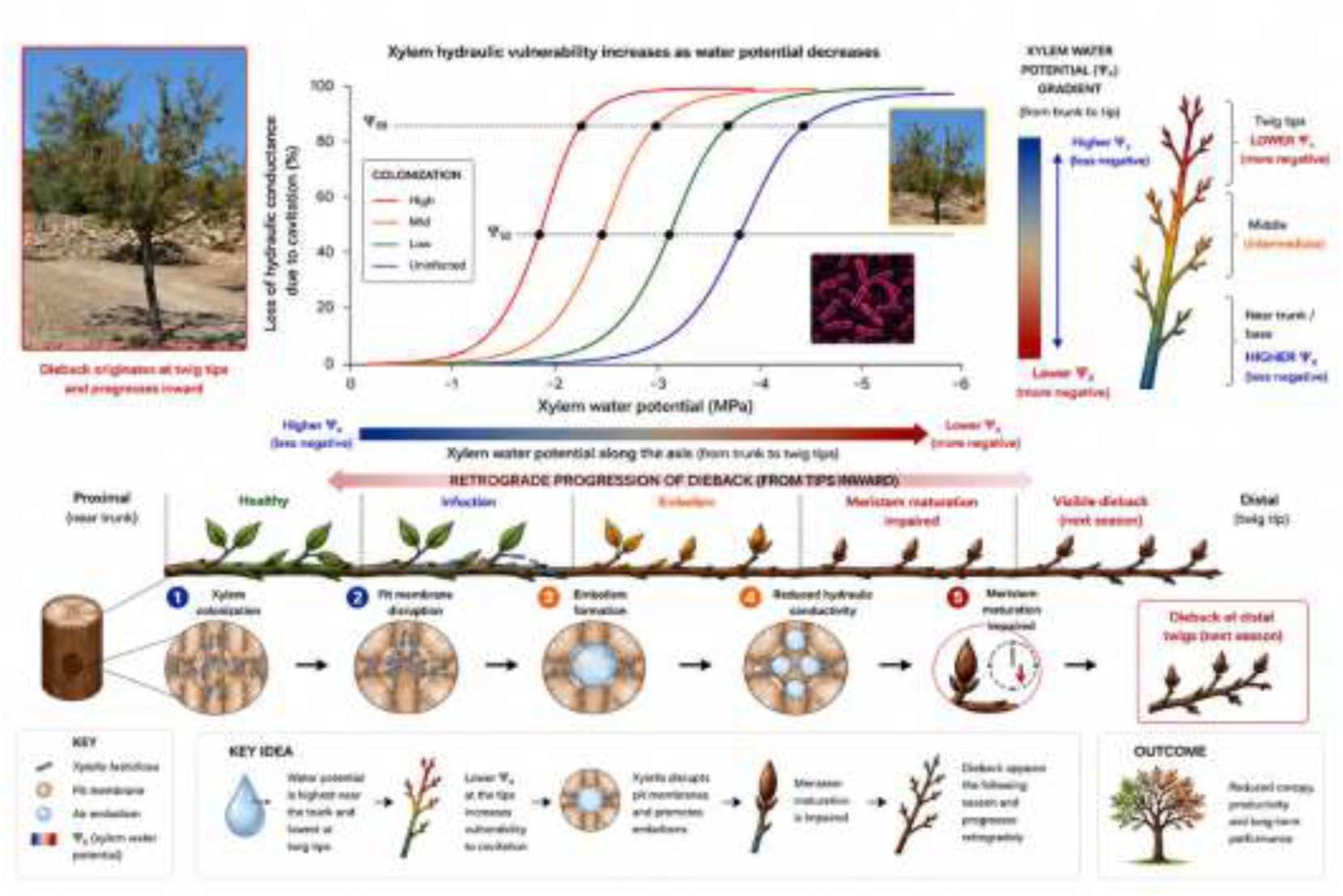
Conceptual model linking xylem water potential gradients, hydraulic vulnerability and retrograde dieback caused by *Xylella fastidiosa* in almond twigs. The figure illustrates a proposed mechanism underlying distal twig dieback in almond trees infected by *X. fastidiosa*. Xylem water potential (Ψ_x_) becomes progressively lower (more negative) from the trunk toward distal twig tips, increasing hydraulic vulnerability to cavitation and embolism formation. The hydraulic vulnerability curves show that increasing bacterial colonization shifts cavitation resistance, leading to conductivity loss at less negative water potentials. Distal tissues, which operate closer to hydraulic failure thresholds, are therefore more susceptible to dysfunction. The proposed sequence involves bacterial colonization of xylem vessels, disruption of pit membrane function, embolism formation, and reduced hydraulic conductivity, ultimately impairing meristem maturation. Symptoms become visible during the following growing season as retrograde dieback progressing from twig tips inward. The model highlights the interaction between xylem hydraulic architecture and pathogen colonization in shaping symptom development and long-term canopy decline in almond trees.

When tracking lineage-specific records across the host population, no significant differences in host longevity or cumulative mortality risk were detected between pathogen subspecies. In the Cox proportional hazards framework, host colonization by *X. fastidiosa* subsp. *multiplex* did not significantly alter the long-term mortality trajectory relative to infection by subsp. *fastidiosa* (Hazard Ratio = 1.04, 95% CI = 0.22–4.99, *P* = 0.958). Although no statistically significant differences in mortality risk were observed between pathogen lineages, these comparative survival metrics should be interpreted with caution. The limited sample size available for distinct subspecies restricts the statistical power of the time-to-event estimators, preventing definitive conclusions regarding lineage-specific virulence trajectories.

## Discussion

The absence of a sustained reduction in radial growth in *X. fastidiosa*–infected almond trees likely reflects a pronounced temporal decoupling between host vegetative productivity and pathogen proliferation. In Mediterranean climates, leaf development and the primary phase of carbon assimilation occur between February and mid-June—a narrow window that determines earlywood formation and, consequently, annual ring width. Conversely, *X. fastidiosa* populations typically begin to build up later in the season, from mid-May onward, with visible disease symptoms rarely manifesting before late June (Mircetich *et al*., 1976; Moralejo *et al*., 2020). Consequently, the main window of cambial activity largely precedes peak bacterial proliferation, minimizing any immediate, direct imprint of infection on xylem radial investment. This phenological mismatch contrasts sharply with pathosystems where pathogen activity directly synchronizes with early-season photosynthesis—such as powdery mildew in oaks— where immediate foliage damage drastically truncates annual radial growth (Bert *et al*., 2016).

Nevertheless, this weak chronological signal in tree rings does not imply negligible functional degradation. Chronic *X. fastidiosa* colonization compromises xylem hydraulic integrity via pit membrane degradation, occlusion of vessels by biofilms, and increased vulnerability to cavitation (Newman *et al*., 2003; Pérez-Donoso *et al*., 2010; Sabella *et al*., 2019). In Mediterranean environments, such hydraulic weakening may remain sublethal during the spring growth surge but becomes critical during late-summer drought and atmospheric vapour pressure deficit stress, when trees operate close to their hydraulic safety margins (Forner *et al*., 2018).

These delayed impacts are therefore more likely to compromise distal meristem survival and bud maturation than stem radial growth itself, providing a mechanistically plausible explanation for the lagged expression of shoot dieback, branch retrogression, and canopy decline (Fig. 5) despite relatively preserved radial growth in preceding years (Mantova *et al*., 2022).

Our findings support a conceptual shift in how chronic vascular infections are integrated into woody plant ecology. Rather than acting as constant, systemic suppressors of biomass accumulation, xylem-limited pathogens can function as latent biotic stressors that remain hidden under average environmental conditions but become functionally disruptive under climate extremes. Within this framework, chronic infection does not primarily alter baseline performance; instead, it fundamentally reshapes the sensitivity of host physiology to climatic variability. Consistently, while neither binary infection status, bacterial load, nor visual symptom severity explained variations in radial growth, climatic water deficit emerged as a dominant predictor. Crucially, the fact that pathogen subspecies significantly modified growth responses to climate demonstrates that biologically meaningful variation is overlooked when infection is treated as a simple binary factor.

This behaviour aligns with threshold models in plant stress ecology, where performance is maintained via compensatory mechanisms until critical physiological tipping points are approached, beyond which minor additional perturbations trigger disproportionate functional collapse (Anderegg *et al*., 2015a; Rowland *et al*., 2015). Chronic vascular infection effectively erodes hydraulic safety margins in ways that are structurally invisible in annual rings, yet highly consequential for drought vulnerability. Similar cryptic dynamics have been observed in other complex pathosystems, where infected individuals track interannual environmental variability in parallel with non-infected cohorts despite underlying physiological impairment (Sisterson *et al*., 2008).

Although our study focuses on the *X. fastidiosa*–almond model, the underlying ecological mechanisms likely extend to a broader class of xylem-inhabiting microorganisms and woody hosts. Myriads of vascular-associated microbes persist asymptomatically within wild plant tissues and are currently ignored in predictive models of vegetation performance or mortality (Bettenfeld *et al*., 2020). Yet, by subtly modifying hydraulic conductance, carbon allocation, or tissue turnover, these cryptic organisms can systematically bias host responsiveness to climatic variability. This suggests that current frameworks of drought-induced forest decline, which rely almost exclusively on abiotic drivers, underestimate the contribution of pervasive, below-threshold biotic factors.

This phenomenon may also reflect a specific evolutionary mismatch where a vascular microorganism interacts with hosts that lack a shared evolutionary history. In its native Neotropical and North American ranges, where long-term host–microbe coevolution is thought to have occurred, *X. fastidiosa* is believed to persist predominantly as a commensal or weakly pathogenic endophyte with negligible fitness costs under benign conditions (Newman et al., 2004; Almeida & Nunney, 2015). Under this evolutionary perspective, functional impacts emerge primarily during climatic anomalies, when reduced hydraulic safety margins or subtle vessel clogging become ecologically costly. Although the baseline prevalence of *X. fastidiosa* in complex Neotropical forests remains unquantified, this scenario is highly consistent with the remarkably low background mortality observed in tropical canopies even during extreme El Niño droughts (Rowland *et al*., 2015; McDowell *et al*., 2018), suggesting that chronic microbial symbioses frequently remain functionally silent under average climate regimes.

Integrating latent biotic stress into plant–climate frameworks is therefore essential for predicting ecosystem resilience under ongoing global change. As drought frequency and intensity accelerate, previously hidden constraints associated with chronic infections will increasingly manifest, driving non-linear patterns of forest decline and agricultural loss. Recognizing that woody plant performance is systematically modulated by persistent, invisible biological agents challenges the traditional dichotomy between biotic and abiotic stress, calling for a unified framework of plant ecological functioning.

## Acknowledgements

We gratefully acknowledge the Plant Health Laboratory of the Balearic Islands for conducting sample analyses. This research was supported by the BEXYL project. We also thank the Conselleria d’Agricultura for their support.

## Competing interests

The authors declare no competing interests.

## Author contributions

E.M. conceived and designed the study, coordinated the research, performed tree-ring measurements, conducted statistical analyses, and led the writing of the manuscript. M.M. coordinated sampling campaigns, performed tree-ring analyses, and carried out DNA extraction from tree-ring samples. B.B.L. conducted molecular analyses of tree rings, including subspecies detection, contributed to manuscript revision, and secured funding. D.O. performed laboratory molecular analyses, contributed to coordination, revised the manuscript, and secured funding.

## Data availability

Code availability: The complete R workflow used for data processing, statistical analyses, and figure generation is available at: https://github.com/emoralejor/hidden-biotic-stress-climate-sensitivity

## Notes

https://github.com/emoralejor/hidden-biotic-stress-climate-sensitivity

### Competing Interest Statement

The authors have declared no competing interest.

