## Supplementary Figure for "Hidden biotic stress alters climate sensitivity in woody plants"

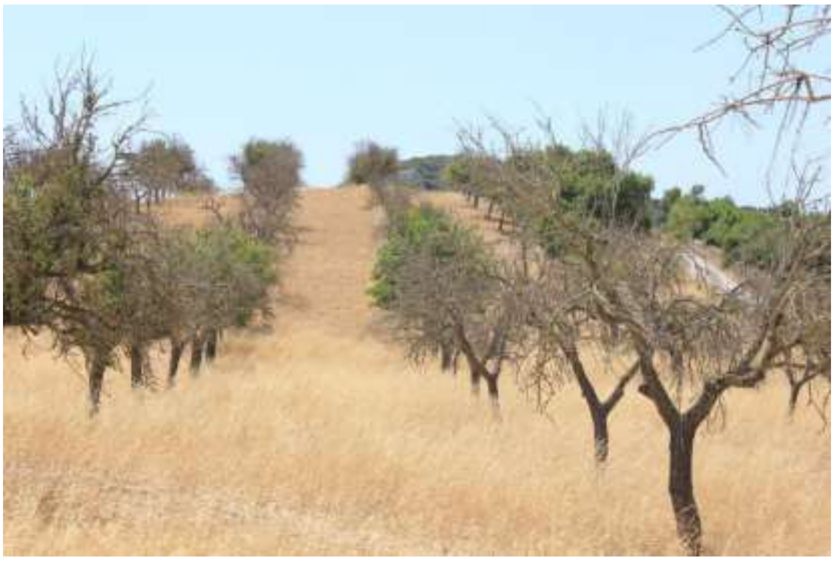

### This PDF file includes:

- **Supplementary note**: Epidemiological context of almond leaf scorch disease in Mallorca
- **Figure S1.** Visual symptomatology scale used for scoring *Xylella fastidiosa* infection severity in almond trees.
- **Figure S2.** Distribution of almond leaf scorch disease severity scores across sampled orchards in 2012.
- **Figure S3.** Seasonal dynamics of symptom severity associated with *Xylella fastidiosa* infection in relatively young almond trees in Mallorca (Spain), monitored from 2017 to 2024.
- **Figure S4.** Variation in almond mortality between 2012 and 2017

**Supplementary Note S1. Epidemiological context of almond leaf scorch disease in Mallorca**

Almond leaf scorch disease (ALSD), caused by *Xylella fastidiosa*, represents one of the largest documented epidemics affecting a perennial crop in Europe. Since its introduction into Mallorca, estimated to have occurred in the early 1990s, the disease has spread across virtually all almond-growing regions of the island. Official surveys conducted since 2017 consistently report widespread pathogen occurrence, with infection prevalences approaching 80–90% in many almond-growing areas. The epidemic has affected more than one million almond trees, making Mallorca a unique natural system for investigating the long-term consequences of chronic vascular infection under Mediterranean climatic conditions.

Current evidence indicates that the epidemic is associated with the coexistence of two *X. fastidiosa* subspecies, *fastidiosa* (ST1) and *multiplex* (ST81), both of which induce a remarkably similar syndrome characterized by progressive leaf scorch, branch dieback, canopy decline, and eventual tree mortality. Although symptom severity varies among individual trees and cultivars, disease progression follows a broadly consistent temporal pattern, with years of apparently normal growth preceding progressive canopy deterioration.

This exceptional epidemiological setting provided the opportunity to investigate long-term host responses using dendrochronology. Rather than attempting to characterize the entire epidemic through exhaustive sampling, our objective was to reconstruct the temporal dynamics of infection and growth in a representative subset of trees exhibiting the characteristic ALSD decline syndrome. The 30 trees selected for this study were drawn from a much larger population of infected individuals displaying comparable symptom development across different locations on the island. For each sampled tree, infection status, bacterial DNA concentration, pathogen subspecies, and annual growth records were comprehensively characterized, allowing the integration of dendrochronological, pathological, and climatic information at annual resolution.

Although this study focuses on a relatively small number of individuals, the strength of the approach lies in the depth of temporal information recovered from each tree. The resulting chronology encompasses 706 annual growth rings, spanning several decades before and after infection, and provides a detailed reconstruction of host responses to chronic vascular colonization under natural field conditions.

The Mallorca epidemic therefore represents a large-scale natural experiment in which thousands of trees have experienced chronic infection under similar climatic conditions over several decades. This unique combination of epidemic scale, well-characterized pathogen populations, and precisely dated annual growth records provides an exceptional opportunity to disentangle the long-term effects of vascular infection from interannual climatic variability.

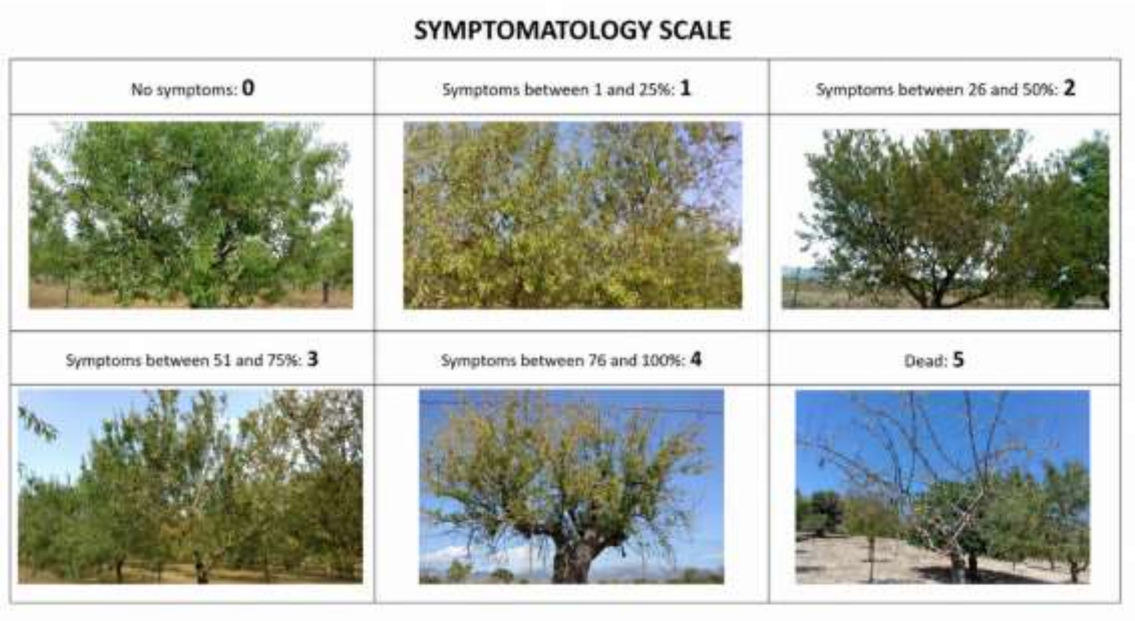


**Figure S1.** Visual symptomatology scale used for scoring *Xylella fastidiosa* infection severity in almond trees. Canopy decline, leaf scorch, and defoliation levels were visually assessed and classified into a 6-class ordinal index (0 to 5): Class 0: asymptomatic canopy; Class 1: 1–25% canopy symptoms; Class 2: 26–50% canopy symptoms; Class 3: 51–75% canopy symptoms; Class 4: 76–100% canopy symptoms with severe dieback; and Class 5: dead tree.

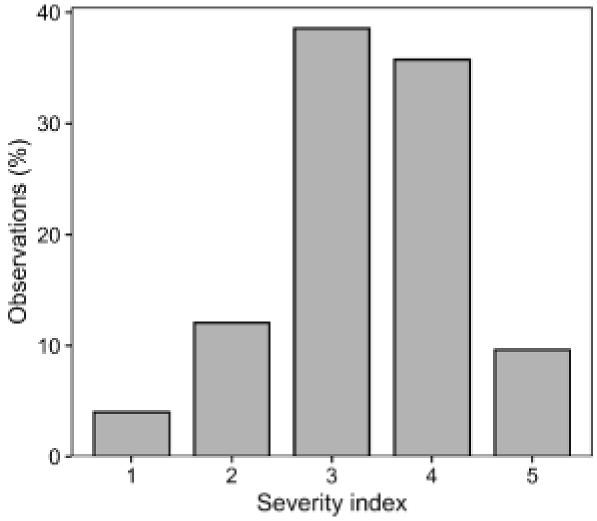


**Figure S2.** Distribution of almond leaf scorch disease severity scores across sampled orchards in 2012. Bars show the percentage of orchard-level observations assigned to each severity class; observations without recorded severity values were excluded.

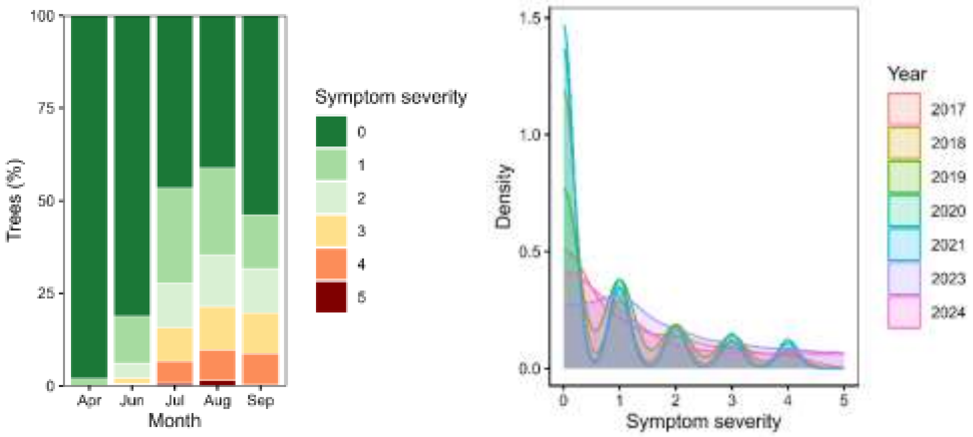


**Figure S3.** Seasonal dynamics of symptom severity associated with *Xylella fastidiosa* infection in relatively young almond trees in Mallorca (Spain), monitored from 2017 to 2024. Stacked bars represent the monthly proportion (%) of individuals within each symptom severity class (0–5). A consistent intra-annual pattern is observed across years, with symptom expression. B, temporal changes in symptom severity associated with *Xylella fastidiosa* in almond trees in Mallorca (Spain) from 2017 to 2024. Density curves (smoothed representations) illustrate the distribution of discrete severity scores (0–5) for each year. The progressive shift towards higher values indicates increasing disease severity over time. increasing during the growing season and declining thereafter, indicating a recurrent seasonal cycle of disease development under Mediterranean conditions.

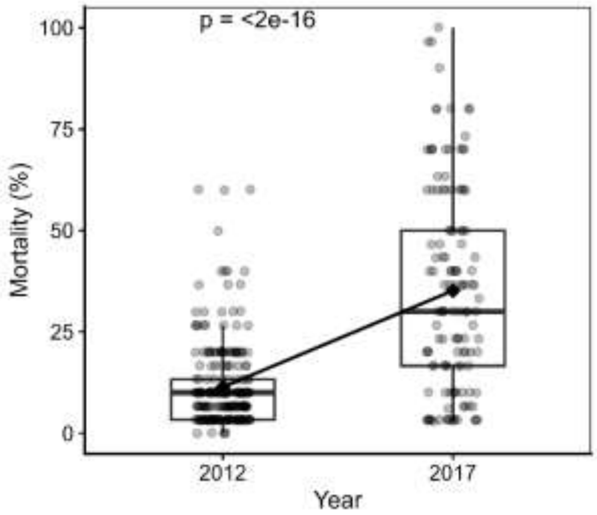


**Figure S4.** Variation in almond mortality between 2012 and 2017. Boxes denote interquartile ranges, horizontal lines indicate medians, and diamonds represent mean values. Points correspond to orchard-level observations, each derived from counts of 30 trees surveyed from a fixed observation point.
